# The macaque ventral intraparietal functional connectivity patterns reveal an anterio-posterior specialization mirroring that described in human ventral intraparietal area

**DOI:** 10.1101/2024.09.20.614031

**Authors:** Wei-An Sheng, Simon Clavagnier, Mathilda Froesel, Wim Vanduffel, Tobias Heed, Suliann Ben Hamed

**Affiliations:** Institut des Sciences Cognitives Marc Jeannerod, UMR5229, CNRS-University of Lyon 1, France; Leuven Brain Institute KU Leuven Medical School, Leuven, 3000, Belgium; Department of Psychology and Centre for Cognitive Neuroscience, University of Salzburg, Salzburg, Austria

**Keywords:** Parietal cortex, ventral intraparietal area, macaque monkey, functional connectivity, resting-state, functional gradient, functional specialization, fMRI, cross-species

## Abstract

The macaque monkey’s ventral intraparietal area (VIP) in the intraparietal sulcus (IPS) responds to visual, vestibular, tactile and auditory signals and is involved in higher cognitive functions including the processing of peripersonal space. In humans, VIP appears to have expanded into three functionally distinct regions. Macaque VIP has been divided cytoarchitonically into medial and lateral parts; however, no functional specialization has so far been associated with this anatomical division. Functional MRI suggests a functional gradient along the anterior-posterior axis of the macaque IPS: anterior VIP shows visio-tactile properties and face preference, whereas posterior VIP responds to large-field visual dynamic stimuli. This functional distinction matches with functional differences among the three human VIP regions, suggesting that a regional specialization may also exist within macaque VIP. Here, we characterized the ipsilateral, whole-brain functional connectivity, assessed during awake resting state, along VIP’s anterior-posterior axis by dividing VIP into three regions of interest (ROIs). The functional connectivity profiles of the three VIP ROIs resembled anatomical connectivity profiles obtained by chemical tracing. Anterior VIP was functionally connected to regions associated with motor, tactile, and proprioceptive processing and with regions involved in reaching, grasping, and processing peripersonal space. Posterior VIP had the strongest functional connectivity to regions involved in motion processing and eye movements. These profiles are consistent with the connectivity profiles of the anterior and posterior VIP areas identified in humans. Viewed together, resting state functional connectivity, task-related fMRI and anatomical tracing consistently suggest specific functional specializations of macaque anterior and posterior VIP. This specialization corroborates the distinction of VIP into three anatomically and functionally separate VIP areas in humans.

## Introduction

The macaque ventral intraparietal area (VIP) is one of several anatomically and functionally distinguishable regions in and around the macaque intraparietal sulcus. It is located in the fundus of the intraparietal sulcus (IPS). VIP has been implicated in a diverse range of sensory, sensorimotor and cognitive functions (Duhamel et al., 1997; Bremmer et al., 2002b, 2002a; Avillac et al., 2004, 2007, reviewed in Foster et al., 2022). The region’s cytoarchitecture (Lewis and Van Essen, 2000a) and structural connectivity (Lewis and Van Essen, 2000b) have also been well defined.

Much effort has gone into defining analogue regions of the monkey’s intraparietal regions in the human brain. However, cytoarchitecture data is incomplete (Amunts et al., 2020; Niu et al., 2020) and matching functional activations to anatomical regions is not straight-forward. One strategy to delineate human functional areas is to assume homology between human and macaque brain areas when interpreting human fMRI data (Mantini et al., 2012; Vanduffel et al., 2014). This strategy has led to suggestions of human homologues of the macaque anterior intraparietal area (AIP), (Binkofski et al., 1999; Culham et al., 2003; Frey et al., 2005; Cavina-Pratesi et al., 2007; Mantini et al., 2012), area V6A (Pitzalis et al., 2013, 2015), and the lateral intraparietal area (LIP), (Denys et al., 2004; Konen and Kastner, 2008). In the case of LIP, however, it appears that macaque LIP has diversified into several neighbouring but distinct regions in humans (Kastner et al., 2017). Regarding ventral intraparietal area (VIP), several incompatible suggestions have been made for a human homologue region, with currently no consensus.

We have recently reviewed the literature about macaque and human VIP and found that fMRI based human VIP studies often use one of three distinct approaches to localize human VIP for further characterization (Foster et al., 2022). These three approaches are (#1) egomotion-consistent visual motion as opposed to local motion (Wall and Smith, 2008), (#2) conjunction of neural responses to visual, tactile, and auditory motion stimuli (Bremmer et al., 2001), and (#3) topographic mapping of tactile and visual stimuli to the face (Sereno and Huang, 2006). Each approach focuses on a subset of macaque VIP functional responses and has consistently resulted in identification of three different locations for a putative human VIP. Most locations that other human fMRI studies have interpreted as the “human VIP” cluster around these three locations. Our review revealed that these three clusters exhibit distinct functional properties and we therefore proposed that the human homologue of macaque VIP is a complex of three distinct regions hVIP#1, hVIP#2 and hVIP#3 (with h denoting “human”); thus, just like LIP, VIP appears to have diversified and expanded in the human brain, as compared to the macaque monkey’s singular region.

The most posterior cluster hVIP#1 has a border with LIP and was reported to be active in studies that used large field optic flow and required processing of self-motion. The most lateral cluster hVIP#2 borders with AIP and was activated in studies that applied neck-proprioceptive stimulation and employed tasks that involved dissociating self-and object-motion. The most anterior cluster hVIP#3 contains topographically organized, tactile and visual face maps, and activations were observed in the context of active head movements.

In principle, as suggested thus far, the three hVIP areas could form one large area with different functional specializations, rather than three distinct areas. Whether a brain region should be classified as distinct depends on multiple criteria, however, such as cytoarchitectonics, connectivity profile, topographic organization (when present), and functional properties (Orban et al., 2004; Orban, 2016). Each hVIP cluster falls into distinct cytoarchitectonic region(s) of published cytoarchitectonic parcellations (Scheperjans et al., 2008; Richter et al., 2019; Amunts et al., 2020). The three hVIPs were distinct also in a parcellation based on anatomical connectivity assessed with diffusion-weighted tensor imaging (DTI) (Mars et al., 2011). In addition, phase-encoded mapping in human PPC revealed retinotopic, saccade-related maps in the location of hVIP#1 (Konen and Kastner, 2008), retinotopic, smooth pursuit-related maps in the location of hVIP#2 (Konen et al., 2005), and somatotopic maps in the location of hVIP#3 (Huang et al., 2017). Thus, each of the three hVIPs appears topographically organized, though for different stimulus characteristics. Viewed together, anatomical and functional characteristics of the three VIPs are most consistent with the view that they are three distinct areas.

This proposal raises the question whether macaque area VIP is truly a single area, or whether it, too, can be subdivided into multiple regions. In fact, cytoarchitectonic mapping divides macaque VIP into a medial and a lateral part (Lewis and Van Essen, 2000a) but no differences in connectivity or physiological functions have so far been associated with this division (Lewis and Van Essen, 2000b). In contrast, functional fMRI studies in macaques have revealed functional differences along its anterior-posterior axis, with anterior VIP responding preferentially to visuo-tactile stimuli around and on the face, and posterior VIP responding preferentially to large-field motion (Guipponi et al., 2013, 2015; Wardak et al., 2016). Here, we used functional connectivity as another means to address potential regional specialization by characterizing resting-state fMRI connectivity of macaque VIP (Mars et al., 2018; Vijayakumar et al., 2018; Gamberini et al., 2020). Specifically, we divided VIP into three regions of interest (ROI) along its anterior-posterior axis and characterized the ipsilateral whole-brain connectivity of these three seed ROIs. The functional networks into which each of the three ROIs was embedded were clearly distinct, in agreement with their respective functions. Thus, functional resting state connectivity profiles support the proposal that macaque VIP exhibits an anterior-posterior functional gradient. Moreover, comparison of the functional division within macaque VIP with the functions ascribed to the three human hVIPs suggests that the distribution of function in the macaque may correspond to the areal diversification observed in the human brain.

## Material and Methods

We reused data from previous studies of Wim Vanduffel’s lab (Mantini et al., 2011, 2013; Pijnenburg et al., 2019). These data were produced in different experiments, however, the recording parameters were similar regarding the puposes and analyses of the present paper.

### 1. Subjects

10 rhesus monkeys (Macaca mulatta) participated in the study (6 females, 4 males). They sat in a sphinx position inside a plastic primate chair (Vanduffel et al., 2001), with their heads constrained by a plastic headpost. The monkeys were trained to maintain fixation at a small red fixation point on a uniform gray background within a 2° × 3° virtual window in the center of the screen, while eye movements were monitored by a pupil-corneal reflection eye tracking system (ISCAN, RK-726PCI) at 60 Hz. The monkeys were rewarded for correct eye fixation behavior, reward frequency increasing to a threshold for longer fixations. Before each scanning session, 8-11 mg/kg monocrystalline iron oxide nanoparticle (Molday ION, BioPAL) was injected via the femoral/saphenous vein to improve the contrast-to-noise ratio (CNR) and to avoid the contribution of superficial draining veins (Vanduffel et al., 2001; Leite et al., 2002). Animal care and experimental procedures were performed in accordance with the National Institute of Health’s Guide for the Care and Use of Laboratory Animal, the European legislation (Directive 2010/63/EU) and were approved by the Animal Ethics Committee of the KU Leuven.

### 2. (f)MRI acquisition

In all studies that produced the data re-used here, in-vivo MRI scans were performed with a 3T MR Siemens Trio scanner in Leuven, Belgium. For functional measurements, gradient echo planar images (GE EPI) covering the whole brain were acquired with an eight-channel phased-array receive coil and a saddle-shaped, radial transmit-only surface coil (see Kolster et al., 2014; 40 slices; 84-by-84 in-plane; flip angle=75°; repetition time (TR) = 2.0 s or 1.4s depending on the monkey; echo time (TE) = 19 ms; voxel size = 1.25 mm x 1.25 mm x 1.25 mm voxels). Specific parameters varied across monkeys (Table 1). T1-weighted anatomical images were obtained during different sessions using a magnetization-prepared rapid gradient echo (MP-RAGE) sequence (TR = 2200 ms, TE = 4.06 ms, voxel size = 0.5 mm x 0.5 mm x 0.5 mm). During the anatomical scans, the animals were sedated using ketamine/xylazine (ketamine 10 mg/kg I.M., xylazine 0.5 mg/kg I.M., maintenance dose of 0.01-0.05 mg ketamine per minute I.V.). The specific acquisition parameters varied across studies and animals (longer runs and higher number of runs were acquired for monkeys achieving better fixation). This resulted in a more robust evaluation of functional connectivity in these individuals. The shortest cumulated resting-state scanning duration (81 min), the shortest duration of individual runs (10 min), and the lowest number of runs (8 runs) combined is in the standard recommendations in the field (Birn et al., 2013). The variability induced by the differences in scanning parameters is thus expected to be minimal, and if anything, it is expected to enhance the robustness of our results, as our approach highlights common connectivity patterns across all monkeys, irrespective of low-level methodological differences.

## 3. Data analysis

### 3.1 Preprocessing

Functional time-series were realigned, coregistered to the anatomical images and normalized to F99 macaque brain template with 1mm isotropic resolution (Van Essen, 2004). The time-series were band-pass filtered (0.01–0.1 Hz or 0.0025 and 0.05 Hz), nuisance corrected for ventricle and white matter signal and motion-scrubbed for movement artefacts within and across sessions and slice time, both by means of regression analysis (Vincent et al., 2007). A spatial smoothing was then applied with a 1.5mm Full Width at Half Maximum (FWHM) Gaussian Kernel.

### 3.2 Region of interest (ROI)

We selected voxels along the IPS by hand from consecutive coronal slices in the volume space of the F99 T1 anatomy for each of the anterior, middle and posterior VIP (aVIP, mVIP and pVIP, see illustration in Fig. 1). For aVIP and mVIP, we selected five voxels each from four consecutive coronal slices. For pVIP, we selected four voxels each from five consecutive coronal slices. These arbitrary subdvisions resulted in all three VIP regions of interest (also later reffered to as sectors) to contain 20 voxels. These voxels were selected using a definition of VIP as provided by the Markov atlas (Markov et al., 2014). The difference in the number of slices assigned to the different ROIs is due to the variation in the dorso-ventral extent of area VIP along the antero-posterior orientation (Fig. 1).

**Figure 1.**
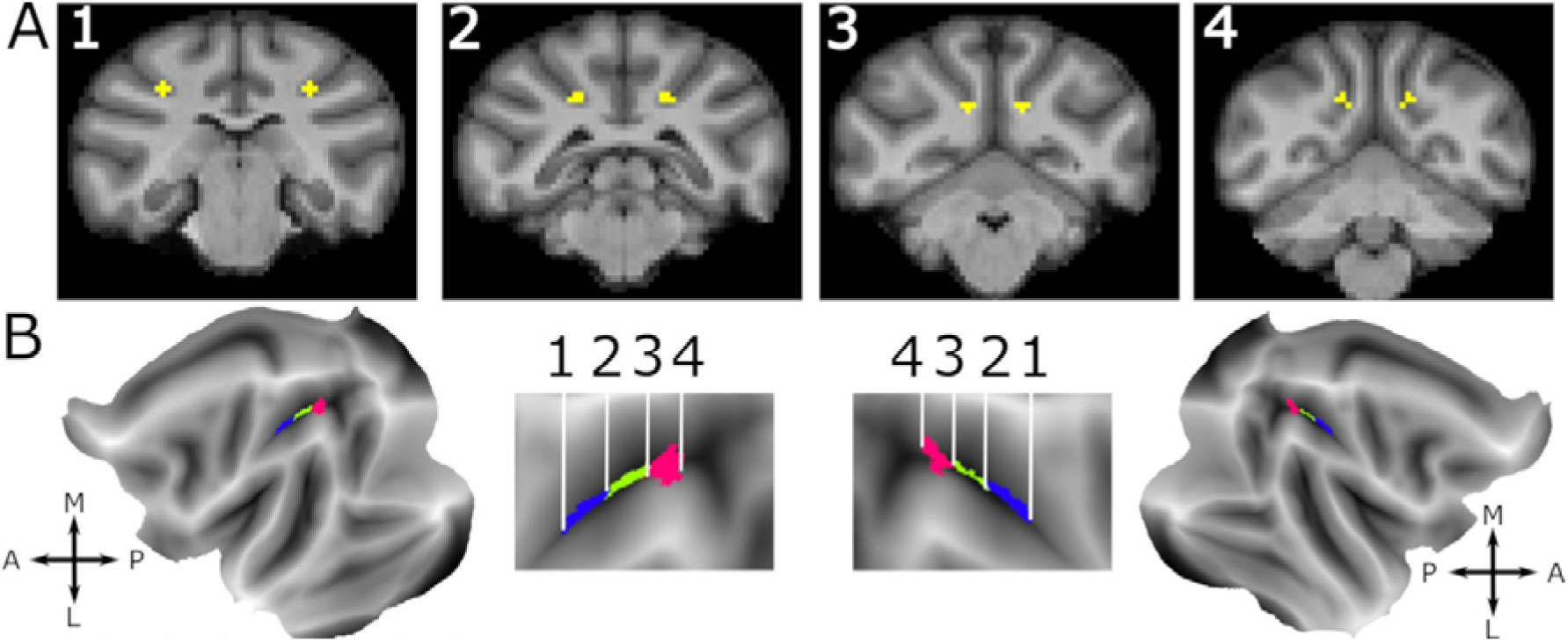
Fundal IPS VIP seed definition in the. F99 T1 anatomy. A. Four example coronal slices in the F99 volume space showing the manually selected voxels in the intraparietal sulcus (IPS) fundus at different antero-posterior levels numbered from 1 to 4. Number 1 and 2 mark the limits of the anterior VIP aVIP ROI. Number 3 and 4 mark the limits of the posterior VIP pVIP ROI. B. The three VIP subregions are projected onto the surface of cortical flat maps for both left and right hemispheres (leftmost and rightmost panel). Middle panel show a zoom on the IPS. The anterior, middle and posterior VIPs are respectively colored in blue, green and pink. Numbers correspond to the section number in A. A, anterior; P, posterior; M, medial; L, lateral, represent brain orientation.

### 3.3 Functional connectivity

Analyses were analyzed using AFNI (Cox, 1996) and FSL (FSL, RID: birnlex_2067; Jenkinson et al., 2012 http://fsl.fmrib.ox.ac.uk/fsl/fslwiki/) embedded in MATLAB. The hemodynamic signal of the 20 voxels of each of the three VIP ROIs was used as seed signal for a seed-to-whole-brain connectivity analysis. Connectivity was calculated as separate correlations of the seed signal and each other voxel of the animal’s brain for each run of each monkey. A Fisher’s r-to-z transformation was applied to the obtained correlation matrix. We then performed, using AFNI, a one sample t-test (against zero) on the resulting maps of all the runs for each ROI, separately for each monkey. This procedure, thus, resulted in one correlation map (z-score) for each VIP ROI and each monkey. We thresholded these maps at p<0.001 (Froesel et al., 2024).

We first projected each individual functional connectivity map onto the brain surface using the Connectome Workbench (Marcus et al., 2013), thus projecting the data from the volume space onto the surface space. Due to the specific mathematical properties of this process, each voxel in the volume space was projected onto one or multiple surface voxels (or faces) on the 2D flatmap (Fig. 2, Fig. 3, Fig. 4, average maps; supplementary Fig. S1, individual monkey maps). We then quantified, across all individuals, the percentage of surface voxels with signal (Fig. 5) and the average z-score of these surface voxels (supplementary Fig. S4), in each of the different functional brain areas as defined by the Markov atlas (Markov et al., 2014).

**Figure 2.**
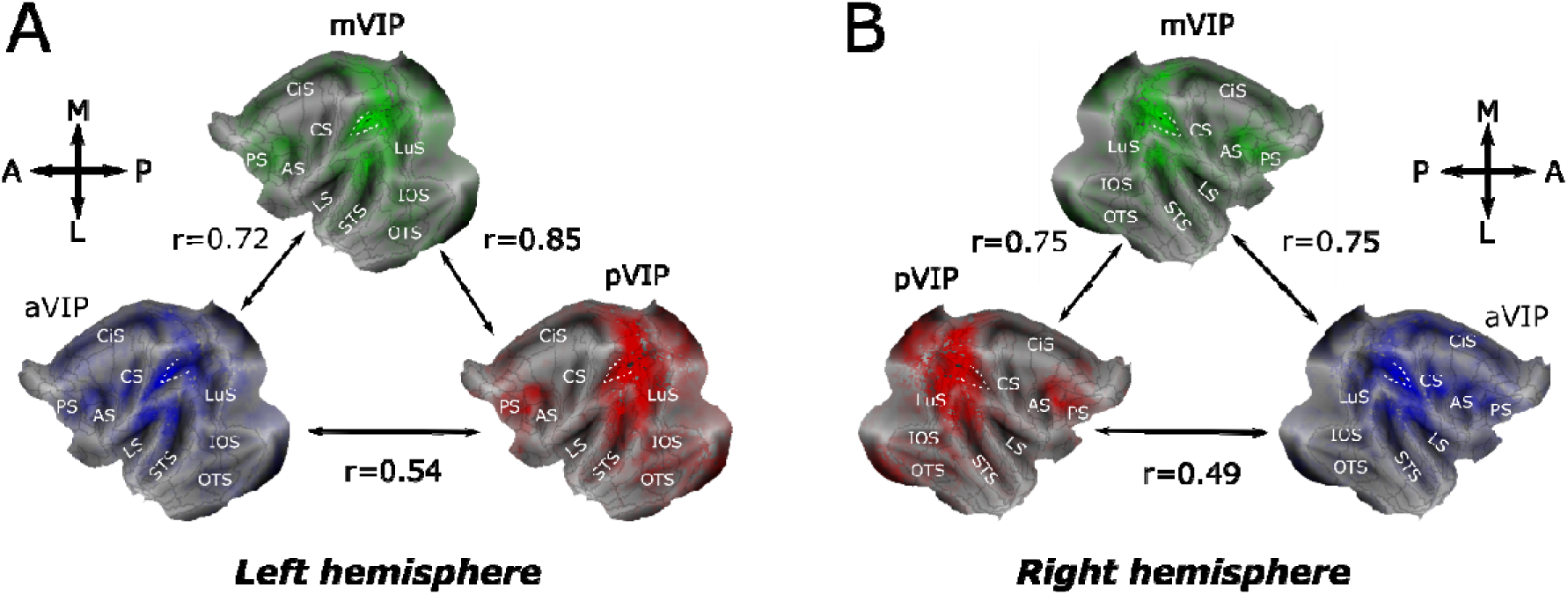
Ipsilateral seed-to-whole-brain functional connectivity patterns for pVIP, mVIP and aVIP regions of interest. Average seed-to-whole-brain ipsilateral functional connectivity patterns of aVIP (blue), mVIP (green) and pVIP (red) ROIs for the left (A) and right (B) hemispheres. Z-score correlation maps from 10 monkeys are averaged and the opacity shows the average connectivity strength on flat maps. Black outlines correspond to the M132 parcellation (Markov et al., 2014). Spearman’s correlation coefficients between pairs of functional connectivity patterns reflect degree of similarity between these patterns (all correlation coefficients have p values <0.05). The IPS is indicated by a white dotted line. AS, arcuate sulcus; CiS, cingulate sulcus; CS, central sulcus; IOS, inferior occipital sulcus; LS, lateral sulcus; LuS, lunate sulcus; OTS, occipitotemporal sulcus; PS, precentral sulcus; STS, superior temporal sulcus.

**Figure 3.**
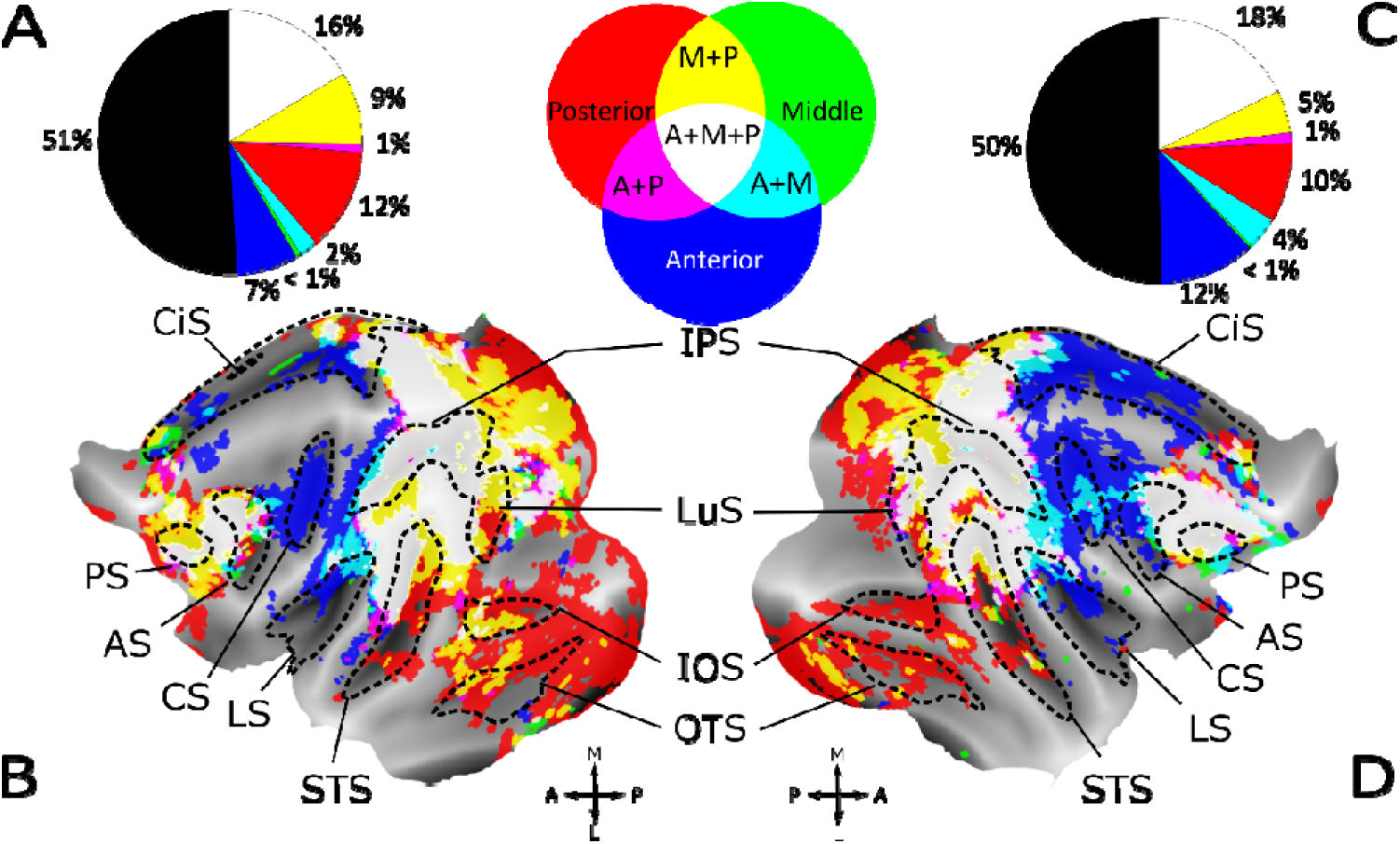
Topographical organization of the cortical regions that are uniquely functionally connected to one of aVIP, mVIP and pVIP ROIs, or to two or more of these three ROIs. Flatmaps of both hemispheres (Figure 3B and D) show regions which are either exclusively connected with the three VIP sectors, or which have joined functional connectivity profiles. Sulci are plotted in dotted lines. Surface voxels with z-scores above 0.05 are colored. Unique, non-overlapping surface voxels functionally connected to the anterior, middle and posterior VIPs are colored in blue, green and red, respectively. The anterior-middle, middle-posterior and anterior-posterior overlapping surface voxels are coded in cyan, yellow and magenta, respectively (middle inset of upper row). Regions functionally connected to all three sectors are shown in white. Pie charts (Figure 3A and C)) illustrate percentages of different categories of unique and overlapping surface voxels in the left and the right hemispheres with the same color code as in the flat maps. Black color in pie charts represents percentage of surface voxels without significant functional connectivity to any of the VIP sectors. AS, arcuate sulcus; CiS, cingulate sulcus; CS, central sulcus; IOS, inferior occipital sulcus; IPS, intraparietal sulcus; LS, lateral sulcus; LuS, lunate sulcus; OTS, occipitotemporal sulcus; PS, precentral sulcus; STS, superior temporal sulcus.

**Figure 4.**
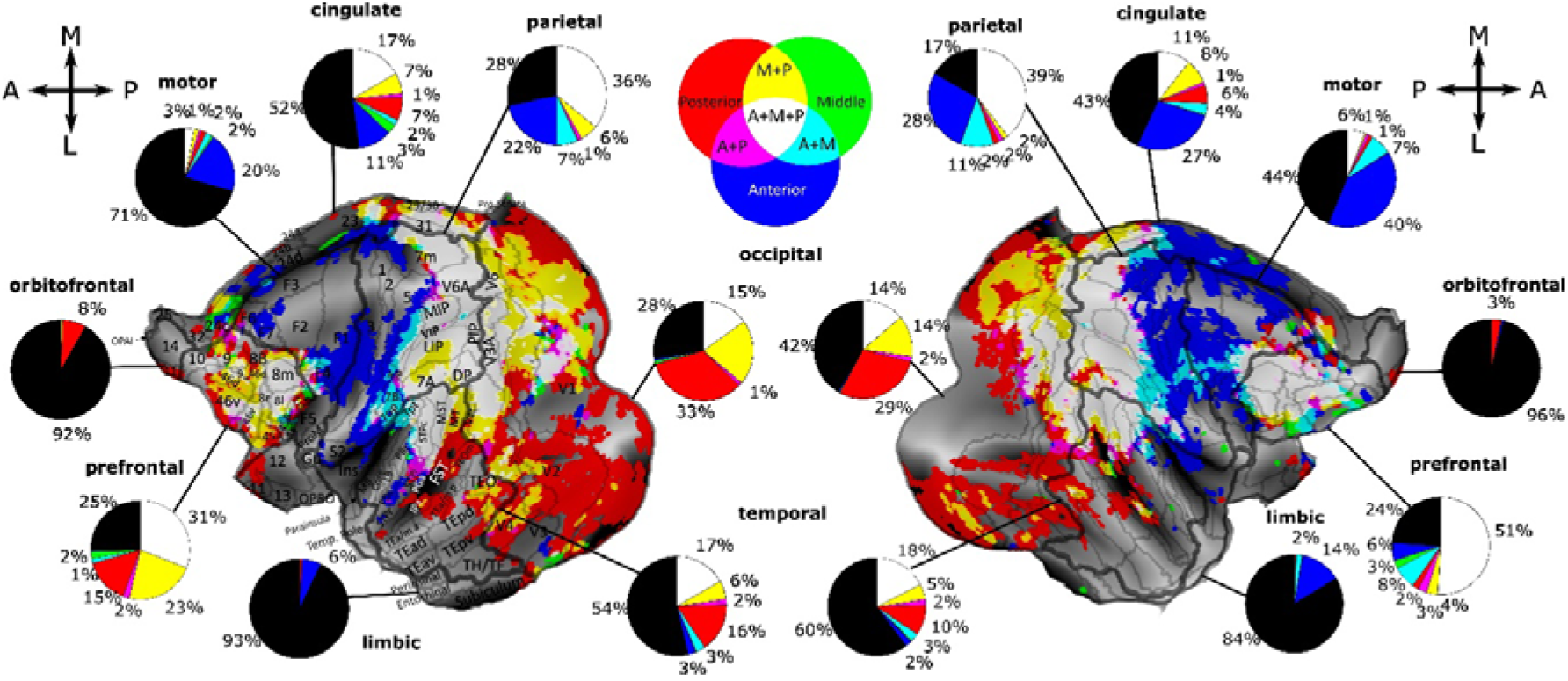
Distributions of the surface voxels in the anterior, middle and posterior ventral intraparietal areas (VIP) that show significant fiunctional connectivity to different parts of the cortex. Flatmaps of both hemispheres show functional connectivity unique to or shared between aVIP, mVIP and pVIP. Surface voxels with z-scores above 0.05 are colored. Unique, non-overlapping surface voxels connected to aVIP, mVIP and pVIP are colored in blue, green and red, respectively. Surface voxels functionally connected to aVIP and mVIP, mVIP and pVIP and aVIP and pVIP are in cyan, yellow and magenta respectively (upper middle inset). Surface voxels functionally connected to all of aVIP, mVIP and pVIP are coded in white. Pie charts show percentages of different categories of unique and overlapping surface voxels in the orbitofrontal, prefrontal, motor/premotor, cingulate, limbic, temporal, parietal and occipital regions. Black color represents percentage of surface voxels lacking significant functional connectivity with VIP. Numerical labels <1% in pie charts are not shown. Areal borders are marked in both hemispheres (Markov et al., 2014), while names are only marked in the left hemisphere. Areas are: 7op, area 7op (parietal operculum); 8l, 8m and 8r, lateral part, medial part and rostral part of area 8; 9/46d and 9/46v, area 9/46 dorsal part and ventral part; 46d and 46v, area 46 dorsal part and ventral part; AIP, anterior intraparietal area, Core, core region of the auditory cortex; DP, dorsal prelunate area; FST, fundus of superior temporal area; Gu, gustatory cortex; LB, lateral belt; LIP, lateral intraparietal area; MB, medial belt; MIP, medial intraparietal area; MST, media superior temporal area; MT, middle temporal area; OPAI, orbital periallocortex; OPRO, orbital proisocortex; PBc and PBr, parabelt caudal part and rostral part; PIP, posterior intraparietal area; S2, secondary somatosensory area; STPc, STPi and STPr, superior temporal caudal part, intermediate part and rostral part; TEad, TEa/ma, TEa/mp, TEav, TEpd and TEpv, area TE, anterior-dorsal part, anterior part, posterior part, anterior-ventral part, posterior-dorsal part and posterior-ventral part; TEOm, area TEO medial part; Tpt, temporo-parietal area; V4t, transitional visual area 4.

**Fig. 5.**
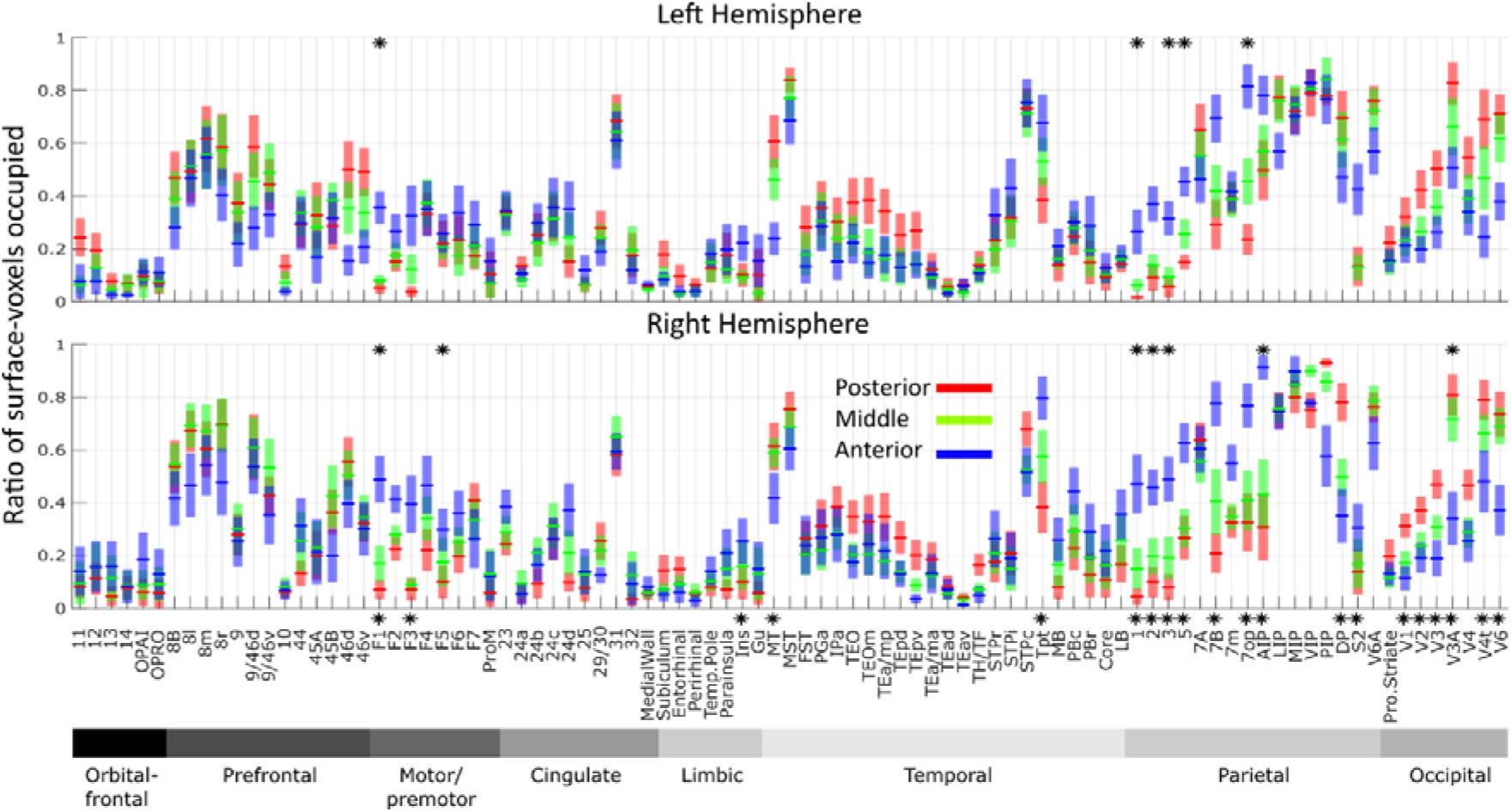
Ratios of functional correlation between aVIP, mVIP and pVIP in atlas-defined cortical functional regions and areas. For each area (Markov atlas, Markov et al., 2014), ratios of surface voxels correlated to aVIP (blue), mVIP (green) and pVIP (red) with z-scores above 0.05 are plotted as line charts for the right and the left hemispheres separately. Shadows show standard error calculated across animals. Areas for which the ratios of surface voxels that are functionally connected to the three VIP seeds in separation are significantly different (Kruskal-Wallis test with Benjamini-Hochberg correction for multiple comparisons, p<0.05) are marked with a star. Tests are performed either on each hemisphere separately (stars on top of the plots), or on both hemispheres (stars at the bottom of the two plots). The areas are grouped by brain regions in the following order: orbitofrontal, prefrontal, motor/premotor, cingulate, limbic, temporal, parietal and occipital. This arrangement orders most areas continually in space with their previous and following areas in the figure. 7op, area 7op (parietal operculum); 8l, 8m and 8r, lateral part, medial part and rostral part of area 8; 9/46d and 9/46v, area 9/46 dorsal part and ventral part; 46d and 46v, area 46 dorsal part and ventral part; AIP, anterior intraparietal area, Core, core region of the auditory cortex; DP, dorsal prelunate area; FST, fundus of superior temporal area; Gu, gustatory cortex; LB, lateral belt; LIP, lateral intraparietal area; MB, medial belt; MIP, medial intraparietal area; MST, media superior temporal area; MT, middle temporal area; OPAI, orbital periallocortex; OPRO, orbital proisocortex; PBc and PBr, parabelt caudal part and rostral part; PIP, posterior intraparietal area; S2, secondary somatosensory area; STPc, STPi and STPr, superior temporal caudal part, intermediate part and rostral part; TEad, TEa/ma, TEa/mp, TEav, TEpd and TEpv, area TE, anterior-dorsal part, anterior part, posterior part, anterior-ventral part, posterior-dorsal part and posterior-ventral part; TEOm, area TEO medial part; Tpt, temporo-parietal area; V4t, transitional visual area 4.

For visualization, we selected surface voxels with a z-score above 0.05 (see supplementary Fig. 3 for results with different thresholds). For Figs. 3, 4, and supplementary Fig. 1, we display the three VIP ROI’s maps on a single surface; non-overlapping surface voxels from the aVIP, mVIP and pVIP were colored in blue, green and red, respectively. Overlap between them is displayed by adding the RGB color values of those colors that overlap, resulting in cyan coloring for anterior-middle VIP overlap, yellow for middle-posterior overlap and magenta for anterior-posterior overlap. Surface voxels containing signals from all three maps are depicted in white.

Last, we assessed the similarity of the three ROIs’ whole-brain functional connectivity maps by a Spearman’s correlation analysis. Fig. 2 illustrates the similarity obtained for the whole-brain functional connectivity maps averaged across all monkeys. Supplementary Fig. 2 represents the distributions of the Spearman’s correlation coefficients between the ipsilateral aVIP, mVIP and pVIP whole brain connectivity maps across individual monkeys.

## Results

In Foster et al., (2022), we proposed that human VIP has diversified into three functionally distinct regions. We wondered, therefore, whether the functional specialization evident in the connectivity of macaque VIP may be regarded as a precursor of the functional specialization of the human VIP complex. To approach this question, we characterized the functional resting-state connectivity pattern of the anterior, middle, and posterior third of macaque VIP (aVIP, mVIP, and pVIP, respectively) in both hemispheres of 10 macaque monkeys (see Fig. 1). Each of these VIP ROIs served as a seed for a whole-brain functional connectivity analysis.

We describe in detail the functional connectivity of aVIP, mVIP and pVIP with the rest of the brain at different spatial scales, from the whole brain to the level of the cortical area, with the aim of inferring possible functional specificities from these patterns of functional connectivity. Thus, in a first step, we characterized the functional connectivity of aVIP, mVIP and pVIP ROIs with the rest of the cortex and identified the cortical regions that are functionally connected to either the entire VIP, or to only some parts of VIP (Fig. 3). In a second step, we zoomed in to the patterns of VIP functional connectivity with the rest of the brain, and we describe how aVIP, mVIP and pVIP connectivity organizes in the main cortical regions, namely the occipital, the parietal, the temporal, the premotor and moror, the limbic, the cingulate, the prefrontal and the orbitofrontal cortex (Fig. 4). We additionally quantified, for all cortical areas defined by the Markov atlas (Markov et al., 2014), their degree of functional connectivity with aVIP, mVIP and pVIP (Fig. 5). For our report, we focus first on those cortical areas that are dominated by functional connectivity with the anterior part of VIP, that is, aVIP or both aVIP and mVIP. We then shift our attention to the cortical areas dominated by functional connectivity with the posterior part of VIP, that is, pVIP, or both of pVIP and mVIP. We draw extrapolations on the putative functions of aVIP, mVIP and pVIP and parallels to the human VIP areas in the discussion.

### A global antero-posterior gradient of whole brain functional connectivity within area VIP

We first used the aVIP, mVIP and pVIP ROIs to characterize the functional connectivity of the entire brain relative to the antero-posterior axis of VIP. We present averaged mapping results, with individual maps available in Supplementary Fig. 1. Fig. 2 displays the averaged connectivity of the three VIP ROIs in the two hemispheres. It is apparent that the three ROIs share similar connectivity patterns, with strongest connectivity within and around the IPS as well as with the prefrontal cortex anterior to the arcuate sulcus (AS). The connectivity maps of aVIP and mVIP were highly correlated (middle-anterior: left hemisphere r = 0.72, p<0.05, right hemisphere r = 0.75, p<0.05), as were mVIP and pVIP maps (middle-posterior: left hemisphere r = 0.85, p<0.05, right hemisphere r = 0.75, p<0.05). In contrast, the correlation was lower between aVIP and pVIP (left hemisphere: r=0.54, p<0.05, right hemisphere: 0.49, p<0.05). The pattern of similarity between neighboring ROIs with a greater difference between the anterior and posterior sectors is indicative of a gradual modulation of connectivity. Fig. 3 displays the three maps overlayed to allow gauging this gradient more intuitively. Connectivity with anterior regions is dominated by aVIP (Fig. 3, blue), followed by regions connected to aVIP and mVIP (Fig. 3, cyan), and, finally, regions connected to all three ROIs (Fig. 3, white). In analogy, connectivity of posterior regions is dominated by pVIP (Fig. 3, red), followed by regions connected to posterior and medial VIP and, then, regions connected to all the ROIs. This functional anterior-posterior gradient of VIP’s connectivity is in line with prior observations from awake macaque fMRI studies that observed a functional gradient between posterior and anterior VIP (Guipponi et al., 2013, 2015; Wardak et al., 2016).

### Global topographical organization of aVIP, mVIP and pVIP functional connectivity to whole brain

In order to better characterize the functional significance of this antero-posterior aVIP, mVIP and pVIP to whole-brain functional connectivity gradient, we visualized the cortical regions that are connected to all three ROIs, thus highlighting a core network recruiting the entire VIP; those regions that are connected to two of these ROIs; as well as those regions uniquely connected to one of the aVIP, mVIP and pVIP ROIs, thus highlighting their functional specificity. To do so, we set a z-score threshold of 0.05 and we colored the surface voxels using an RGB color code (Fig. 3). When applying this specific threshold of z<0.05, approximately an equal number of cortical surface voxels is either connected to VIP, or not, most probably describing both direct and second-order indirect anatomical connectivity with VIP. Varying this threshold from between 0.03 and 0.07 did not substantially change the observations reported below (see supplementary Fig. S3).

#### Cortical regions functionally connected to all three VIP regions (aVIP, mVIP and pVIP)

In a first step, we focus on the regions that show functional connectivity with all three VIP ROIs. There were two such regions (Fig. 3, white, supplementary Fig. S1 for individual monkeys’ maps). They represent 16% and 18% of the total brain surface in the left and the right hemispheres respectively, and 33% and 36% of VIPs overall functional connectivity to the rest of the cortex. The first region covers a large extent of the parietal cortex and adjacent brain regions. Specifically, it is centered on the intraparietal sulcus (IPS) and extends along a medio-lateral axis. It thus includes the posterior part of the cingulate sulcus (CiS) and medial parietal cortex. It also includes the entire IPS, extending into the inferior parietal cortex up to the lunate sulcus (LuS). Last, it includes the posterior section of the superior temporal sulcus (STS), and extends into the cortical convexity between the STS and the lateral sulcus (LS). The second region is located in the frontal cortex, anterior to the AS and mostly extends within and dorsally to the principal sulcus (PS).

#### Cortical regions functionally connected to only one or two VIP ROIs

In a second step, we focus on the regions that show functional connectivity with only two of our three VIP ROIs (Fig. 3, yellow, cyan and magenta). These represent 12% and 10% of the whole brain surface in the left and the right hemispheres respectively, and 24% and 20% relative to all the cortical surface voxels functionally connected to VIP (pie charts in Fig. 3 for detailed % per category).

The mVIP functional connectivity map exhibits large overlap with pVIP and aVIP functional connectivity maps, likely due to the current resolution of our fMRI data as well as to partial volume effects, resulting in enhanced functional connectivity between near-by voxels. As a result, most cortical surface voxels with significant functional connectivity with mVIP are also significantly functionally connected to either pVIP or aVIP (Fig. 3). We first describe the topographical organization of surface voxels functionally connected to aVIP, or to both aVIP and mVIP. Then, we describe the topographical organization of surface voxels functionally connected to pVIP, or to both pVIP and mVIP.

Surface voxels functionally connected to aVIP and aVIP-mVIP were organized along a gradient (indicated by the white-cyan-blue colors) anterior to the parietal cortex, 1) starting medially from the lateral bank of the CiS, extending along the medial bank of the IPS and ending laterally in the fundus of the LS in both hemispheres; 2) in the central sulcus (CS); as well as 3) in the medial parietal cortex, along the entire extent of the CiS, though with an emphasis on the anterior cingulate cortex. The mVIP-aVIP functionally connected surface voxels (cyan) occupied 2% and 4% of the whole brain surface in the left and the right hemispheres respectively, while functionally connected surface voxels of aVIP (blue) accounted for 7% and 12% of all cortical surface voxels.

Surface voxels functionally connected to pVIP and pVIP-mVIP organize along a gradient (white-yellow-red surface voxels) that extends 1) into the major parts of the occipital cortex, except most of the operculum. Part of this connectivity gradient starts medially at the posterior end of the CiS, and extends along the posterior occipital sulcus (POS) and the posterior bank of the LuS; a second branch starts between the STS and the inferior occipital sulcus (IOS), and occupies a large area in the lateral occipital cortex, including the IOS and the occipitotemporal sulcus (OTS). 2) Furthermore, surface voxels connected to pVIP and pVIP-mVIP are found in the middle of the superior temporal sulcus (STS), as well as 3) around and above the principal sulcus. The surface voxels functionally connected to pVIP-mVIP (yellow) occupy 9% and 5% of the whole brain surface in the left and the right hemispheres respectively, while the pVIP functionally connected surface voxels (red) accounted for 12% and 10% of all cortical surface voxels in both hemispheres, respectively.

Although pVIP and aVIP are non-contiguous VIP sectors, 1% of all cortical surface voxels are connected to these two sectors but not to mVIP between them (magenta). They are located in very small sectors of the temporal, parietal and occipital cortex. Their occurrence cannot be attributed to the current resolution of the functional MRI data. These surface voxels are few relative to the other types of surface voxels. Yet, they are remarkably symmetrical across hemispheres, suggesting that they reflect not mere measurement noise but a consistent organizational pattern across hemispheres and animals. Hence, these surface voxels may contribute to specific cognitive functions which rely on the combination of specific neuronal signals encoded in both pVIP and aVIP.

Last, surface voxels exclusively connected to mVIP (green) occupy less than 1% of the whole brain surface in both hemispheres, and are mainly found in prefrontal and cingulate regions. Again, since they were identified in symmetrical locations in both hemispheres and remotely from the seeding ROIs, i.c., in the medial prefrontal cortex and in the ventral lateral prefrontal cortex, this suggests that mVIP might be exclusively connected to specific cortical areas, not targeted by pVIP and aVIP.

### Cortical regions and areas dominated by functional connectivity to aVIP and aVIP-mVIP

In order to better understand the functional significance of the differences in whole brain functional connectivity of aVIP, mVIP and pVIP, we next assessed their functional connectivity to specific brain regions including orbitofrontal, prefrontal, premotor/motor, cingulate, limbic, temporal, parietal and occipital regions (Fig. 4, same connectivity maps as Fig. 3 but with additional regional and areal broaders), (Markov et al., 2014). We then calculated the percentage of surface voxels functionally connected to aVIP, mVIP and pVIP and their combinations (Fig. 4, pie charts). Additionally, we parcellated the cortex into 92 functionally and anatomically defined cortical areas (Fig. 4), using Markov atlas (Markov et al., 2014). We then calculated the ratios of surface voxels showing significant funcitonal connectivity with each of the individual VIP seeds (Fig. 5) as well as the corresponding average z-scores (supplementary Fig. S4), separately for each region and each area. In the following paragraph, we first discuss the cortical regions expressing a dominance of functional connectivity with aVIP, or aVIP-mVIP. Then, we discuss the cortical regions expressing a dominance of functional connectivity with pVIP, or pVIP-mVIP.

#### Focus on parietal cortex: a posterior-anterior gradient from aVIP-mVIP-pVIP to aVIP-mVIP to aVIP functional connectivity

When we zoom in to connectivity in parietal cortex, the percentage of surface voxels that were functionally connected to all three VIP seeds (white surface voxels) was largest, compared to all other categories (Fig. 4, left hemisphere: 36%, right hemisphere: 39% of all parietal surface voxels; note that in figure 3, percentages are computed over the entire hemisphere while in figure 4, they are computed relative to the specific brain region); this finding is unsurprising given that VIP is located in the parietal cortex. Parietal surface voxels connected to all three VIP ROIs were located in areas LIP, MIP, PIP, 7A and V6A (Fig. 4, Fig. 5, supplementary Fig. 4). The latter areas did not show a preferential functional connectivity pattern to any of the three VIP ROIs (Kruskal-Wallis test, p>0.05).

Twenty-two percent and 28% of the parietal surface voxels in the left and right hemisphere, respectively, were functionally connected to aVIP (Fig. 4, blue surface voxels). Only a small fraction of surface voxels were functionally connected to both aVIP and mVIP (Fig. 4, cyan surface voxels, left hemisphere: 7%, right hemisphere: 7% of all parietal surface voxels). These surface voxels were located in areas 1, 2, 3, 5, 7m, 7op, 7B, AIP, and S2. They were characterized by stronger functional connectivity with aVIP than mVIP and pVIP when pooled across hemispheres (Fig. 5, Kruskal-Wallis test with Benjamini-Hochberg correction, p<0.05).

A yet smaller percentage of parietal surface voxels were functionally connected to aVIP-pVIP (magenta surface voxels), to mVIP-pVIP (yellow surface voxels), or only to pVIP (red surface voxels). These latter types of surface voxels were either very few or located in non-symmetrical location across hemispheres. They were situated in area DP (Markov et al., 2014) also known as Opt (Pandya and Seltzer, 1982) which, overall, had stronger functional connectivity with pVIP than the other two RIOs (Kruskal-Wallis test Benjamini-Hochberg procedure, p<0.05).

Overall, parietal surface voxels thus exhibited a posterior to anterior functional connectivity to aVIP, mVIP and pVIP, such that the most posterior surface voxels were functionally connected to all three VIP ROIs, these were bordered by aVIP-mVIP functionally connected surface voxels, while surface voxels uniquely connected to aVIP were located more anteriorly.

#### Focus on cingulate cortex: a posterior-anterior gradient of aVIP-mVIP-pVIP to aVIP-mVIP to aVIP functional connectivity

Zooming in on cingulate cortex revealed wide-spread functional connectivity to VIP (left hemisphere, 48%; right hemisphere 57%, of all cingulate surface voxels). Posterior cingulate regions were either functionally connected to all three VIP ROIs, to both pVIP-mVIP, or only to pVIP (Fig. 4). This pattern of functional connectivity is very similar to the one we report in the present study in the occipital cortex and mostly concerns area 31 (Fig. 4, Fig. 5). More anteriorly, functional connectivity shifts to aVIP-mVIP and more rostrally to aVIP. This functional connectivity pattern covers areas 23 and 24d and extends dorsally towards the medial part of the premotor cortex (Fig. 4). More anteriorly, functional connectivity is evident with pVIP and pVIP-mVIP bilaterally, and with mVIP in the left hemisphere. This pattern corresponds to a medial extension of the functional connectivity characterizing adjacent prefrontal cortex. Overall, area 31 is the most densely functionally connected cingulate region to the VIP ROIs, followed by areas 24b, 24c, 24d and 23. Areas 25 and 24a, on the other hand are only weakly functionally connected to VIP (Fig. 5, ratio < 20%).

#### Focus on premotor and motor cortex: functional connectivity predominantly to aVIP

Zooming in on the premotor and motor regions reveals a dominance of surface voxels uniquely functionally connected to aVIP in both hemispheres (Fig. 4, left hemisphere, 20%; right hemisphere: 40% of premotor/motor surface voxels). In these regions, F1 to F7 except area ProM were functionally connected to aVIP in both hemispheres (Fig. 5). However, area F1 stood out as an area in which the ratio of surface voxels functionally connected to aVIP were significantly higher than the ratio of surface voxels functionally connected to either mVIP or pVIP in both hemispheres (Kruskal-Wallis test with Benjamini-Hochberg procedure, p<0.05). The other premotor/motor areas had higher ratios of surface voxels functionally connected to pVIP or mVIP. This was particularly marked for areas F4 and F7 (Fig. 4, Fig. 5). In F2, F4 and F5 of both hemispheres, surface voxels connected to mVIP and pVIP form a weak mVIP-aVIP to aVIP (white-cyan-blue) gradient posteriorly to the white surface voxels (i.e. those functionally connected to all three VIP ROIs) centered between the AS and the PS (Fig. 4). In the medial part of the premotor/motor cortex, F3 and F6 were functionally connected mostly to aVIP (Kruskal-Wallis test with Benjamini-Hochberg procedure, p<0.05). In these areas, the ratios of surface voxels correlated with aVIP were higher in the right than in the left hemisphere.

#### Focus on limbic cortex: functional connectivity predominantly to aVIP

Only aVIP shows significant functional connectivity with the insula (relative to mVIP and pVIP when data from both hemispheres are pooled, Kruskal-Wallis test with Benjamini-Hochberg correction, p<0.05, Fig. 5. If the left or right hemispheres are considered separately, both are non-significant). This corresponds to the ventral-most end of the cortex-wide mVIP-aVIP to aVIP (white-cyan-blue) gradient (Fig. 4). Other areas of the limbic system are largely uncorrelated with any of the VIP seeds (Fig. 4, Fig. 5). The aVIP surface voxels make up 6% and 14% of the entire limbic cortex in the left and the right hemispheres, respectively, indicating relatively week functional connectivity of VIP to this region.

### Cortical regions and areas dominated by functional connectivity to pVIP and pVIP-mVIP

#### Focus on occipital cortex: an anterior-to-posterior gradient of aVIP-mVIP-pVIP to pVIP-mVIP to pVIP functional connectivity

The mVIP-pVIP to pVIP (white-yellow-red) gradient posterior to the seed ROIs was mainly located in the occipital and temporal regions (Fig. 4). In the occipital cortex, the pVIP (red) and mVIP-pVIP (yellow) surface voxels occupied 33% and 21% in the left hemisphere, and 29% and 14% in the right hemisphere, respectively (Fig. 4). While functional connectivity to pVIP was predominant, significant functional connectivity to mVIP and to a lesser extent to aVIP was also evident (Fig. 5). Furthermore, ratios of surface voxels that functionally connected to all three VIPs increased from V1 (left: 26%; right: 20%), to V2 (30%, 27%), to V3 (37%, 32%), to V3A (66%, 62%) as well as from V4 (42%, 34%), to V4t (47%, 64%) and to V6 (57%, 60%).

#### Focus on temporal cortex: a medial-to-lateral gradient of aVIP-mVIP-pVIP to pVIP-mVIP to pVIP functional connectivity

Zooming in on temporal cortex revealed that pVIP (red) and mVIP-pVIP (yellow) surface voxels occupy 16% and 6% of the temporal region of the left hemisphere, respectively, and 10% and 5% in the right hemisphere (Fig. 4). The mVIP-pVIP to pVIP (white-yellow-red) global gradient contained two branches, and a small anterior part of the second (lateral) branch was located in the temporal region, especially in TEO and TEOm (Fig. 4). These areas were close to the lateral end of the mVIP-pVIP to pVIP gradient in STS, including FST, PGa and IPa. The connectivity pattern in the aforementioned five areas suggests a weak trend of functional connectivity to pVIP (Fig. 5), but this apparent trend was not statistically significant (Kruskal-Wallis test with Benjamini-Hochberg procedure, p>0.05). The ratio of surface voxels functionally connected to pVIP decreased more laterally in TEam/p and TEpd. Medial to these areas were white surface voxels functionally significantly connected to all three VIP ROIs, including MT, MST, STPc and Tpt (Fig. 4). This was also near the location where the aVIP-mVIP-pVIP to pVIP-mVIP to pVIP gradient and the aVIP-mVIP-pVIP to aVIP-mVIP to aVIP gradient touch each other. As a result, MST and STPc had very high ratios of functionally connected voxels to all three VIPs, while MT and Tpt were dominated by functionally connected surface voxels of pVIP and aVIP, respectively (Kruskal-Wallis test with Benjamini-Hochberg procedure, p<0.05, Fig. 5).

#### Focus on prefrontal cortex: a posterior-to-anterior gradient of aVIP-mVIP-pVIP to pVIP-mVIP to pVIP functional connectivity in the left hemisphere

Zooming in on the prefrontal cortex revealed dense connectivity to all three VIP ROIs, which is remarkable given its remote distance from the IPS. It represented the second largest functionally connected region to all VIP ROIs, covering the AS and the PS and accounting for 51% and 31% of the entire prefrontal region in the right and left hemispheres respectively (Fig. 4). In the left hemisphere, areas 9, 9/46d, 45A, 46d and 46v were strongly functionally connected to pVIP, with weaker functional connectivity to mVIP and aVIP (Fig. 5). In the right hemisphere, functional connectivity was equally strong between these areas and the three VIP ROIs (Fig. 5). In the left hemisphere, an additional mVIP-pVIP to pVIP (white-yellow-red) gradient was evident anteriorly. The pVIP (red) connected surface voxels and the mVIP-pVIP (yellow) surface voxels occupied 15% and 23% of the prefrontal region. In the right hemisphere, there were no obvious mVIP-pVIP to pVIP (white-yellow red) or mVIP-aVIP to aVIP (white-cyan-blue) gradients but a “mosaic composition”, associating functional connectivity from different VIP ROI combination, at the edge of the white surface voxels region. This mosaic composition was larger at the ventral side (Fig. 4). The ratios of the aVIP (6%) and the mVIP-aVIP (8%) connected surface voxels were however higher than the pVIP (2%) and the mVIP-pVIP (4%) surface voxels within the mosaic prefrontal cortex composition. This contrasted with the left hemisphere, where pVIP and mVIP-pVIP surface voxels made up the mVIP-pVIP to pVIP gradient, and where the ratios of the aVIP and the mVIP-aVIP connected surface voxels accounted for less than 2% of the prefrontal region (Fig. 4). Another mosaic organization was evident at the junction of the prefrontal, cingulate and premotor/motor regions, bilaterally, including areas 8B, 9, 24c and F6. In the left hemisphere, this spot was dominated by pVIP (red), mVIP-pVIP (yellow) and aVIP-pVIP (magenta) surface voxels, while the right hemisphere exhibited a mosaic composition (Fig. 4).

#### Focus on orbitofrontal cortex: weak connectivity with VIP, predominantly with pVIP

The connectivity pattern of the orbitofrontal regions resembled that of the limbic regions, with weak functional connectivity with VIP. Indeed, orbitofrontal surfave-voxels showed significant functional connectivity with pVIP, specifically surfave-voxels located in area 11 and the medial portion of area 12 of both hemispheres. The pVIP surface voxels accounted for 8% and 3% in the orbitofrontal regions of the left and right hemispheres (Fig. 4).

## Discussion

We divided macaque VIP into three isovolumetric ROIs along its anterior-posterior axis to identify their whole-brain functional connectivity patterns and to characterize their degree of similarity. Our results confirm previous findings of strong functional connectivity between the fundus of the IPS and the prefrontal cortex anterior to the AS and around the AS from all three ROIs (Lewis and Van Essen, 2000b). In addition, and similarly to what we have described in humans (Foster et al., 2022), whole brain functional connectivity of the posterior part of VIP was different from the whole brain functional connectivity of its anterior part, thus defining an antero-posterior functional gradient in this region. Below, these results are discussed in light of the anatomical and physiological properties of macaque VIP as well as the functional specialization of the putative VIP homologue in humans.

### Comparison of VIP’s anatomical and functional connectivity

Resting-state functional connectivity reflects both anatomical connections (van den Heuvel et al., 2009; Matsui et al., 2011; Wang et al., 2013; Tang et al., 2021; Card and Gharbawie, 2022) and indirect structural connections (Vincent et al., 2007; Honey et al., 2009). In our study, all known VIP anatomical connections (Lewis and Van Essen, 2000b) are also identified in its functional connectivity (Fig. 5). However, we observed additional functional connections that have not been identified anatomically, which thus most probably indicate indirect functional connectivity. A previous tracer study classified VIP connection strength in each cortical area into three levels (Lewis and Van Essen, 2000b, Table 3). The areas in these three different levels corresponded to 34% ± 17% (mean +/-s.d.), 39% ± 19%, and 51% ± 24% surface voxels respectively that are functionally connected to VIP, on average (across areas and hemispheres). While the functional connectivity patterns reported in this studyoverall matched anatomical connectivity strength, the amount of connectivity did not always match between the two types of connectivity. For example, about 10% of surface voxels of area 24a and TEa/ma are functionally connected to VIP (Fig. 5), while 60% of surface voxels of areas 8m and V3A are functionally connected to VIP. Yet both areas 24a and TEa/ma as well as areas 8m and V3A show only weak anatomical connectivity with VIP. Such variability also existed in the group of areas showing the strongest anatomical connectivity with VIP, for example, approximately 20% of F1’s surface voxels are functionally connected to VIP (Fig. 5), while 80% of surface voxels in the posterior intraparietal area (PIP) are functionally connected to VIP. This is in line with a previous study comparing diffusion tensor imaging and resting-state functional connectivity showing that indirect structural connections contribute to unique functional connections without structural connections and introduce variability in functional connection strength (Honey et al., 2009).

Several areas exhibited functional connectivity to VIP although no corresponding anatomical connectivity has so far been reported. Connectivity with orbital-frontal regions was not reported in the seminal anatomical connectivity study by Lewis and Van Essen (2000b, Table 3); yet, in the resting-state data, we found functional connectivity between VIP and area 11 and 12. The extent of this functional connectivity was similar to that observed between VIP and area 24a and TEa/ma (ca. 10% of the surface voxels, Fig. 5), two cortical regions for which weak anatomical connections with VIP have been identified. If we set an arbitrary criterion of 10% of an area’s surface voxels exhibiting connectivity as evidence for functional connectivity, we identified several other cortical brain areas that are functionally connected to area VIP while no previous report of corresponding anatomical connectivity exists. These includes area 9 (including 9/46v and 9/46d from the Markov map) and area 44 in the prefrontal regions, area 24c and area 31 in the cingulate regions, and area V1 in the occipital region; multiple areas in temporal cortex, including area TEO, TEpd and TEpv in area TE, areas TH and TF, as well as the caudal part of the parabelt (PBc), including medial belt, core belt region and lateral belt of the auditory cortex; as well as limbic regions including the temporal pole, parainsula, insula and gustatory cortex.

### Functional homologies between macaque and human VIP subdivisions

The vast number of VIP’s anatomical connections (Lewis and Van Essen, 2000b) can be classified into eight functional categories (Foster et al., 2022) based on their contribution to the functions ascribed to to VIP (Fig. 6). We used these anatomically connected areas and functional networks as a template to test whether the functional specialization of the differential connectivity patterns observed for our three macaque VIP subdivisions match with the three putative VIP homologue clusters we have suggested as the expanded human VIP homologue in the human brain (Foster et al., 2022). Human fMRI studies have applied a specific set of tasks to identify VIP based on a specific function that had previously been reported by research on macaque VIP. We could match the human fMRI task designs associated with each cluster with the eight functions attributed to macaque VIP. According to this matching procedure, the most posterior human VIP homologue area, hVIP#1, is associated with motion detection and eye movements, the more lateral hVIP#2 is associated with proprioception and eye movements, and the most anterior hVIP#3 is associated with tactile processing and peripersonal space representation. To compare the functional connectivity patterns of the VIP subdivisions between humans and macaques, we performed a meta-analysis of the literature and matched the macaque atlas-defined areas to the three human VIP clusters. We then identified to which macaque VIP ROI each macaque area has the strongest correlation with, and compared these results with the human cluster functional network (Fig. 6). As in our analysis no area had significantly stronger conncectivity to the mVIP than to the aVIP or pVIP (Fig. 5 and supplementary Fig. 2), we concentrated our comparison on the latter ROIs (Fig. 6).

**Fig. 6.**
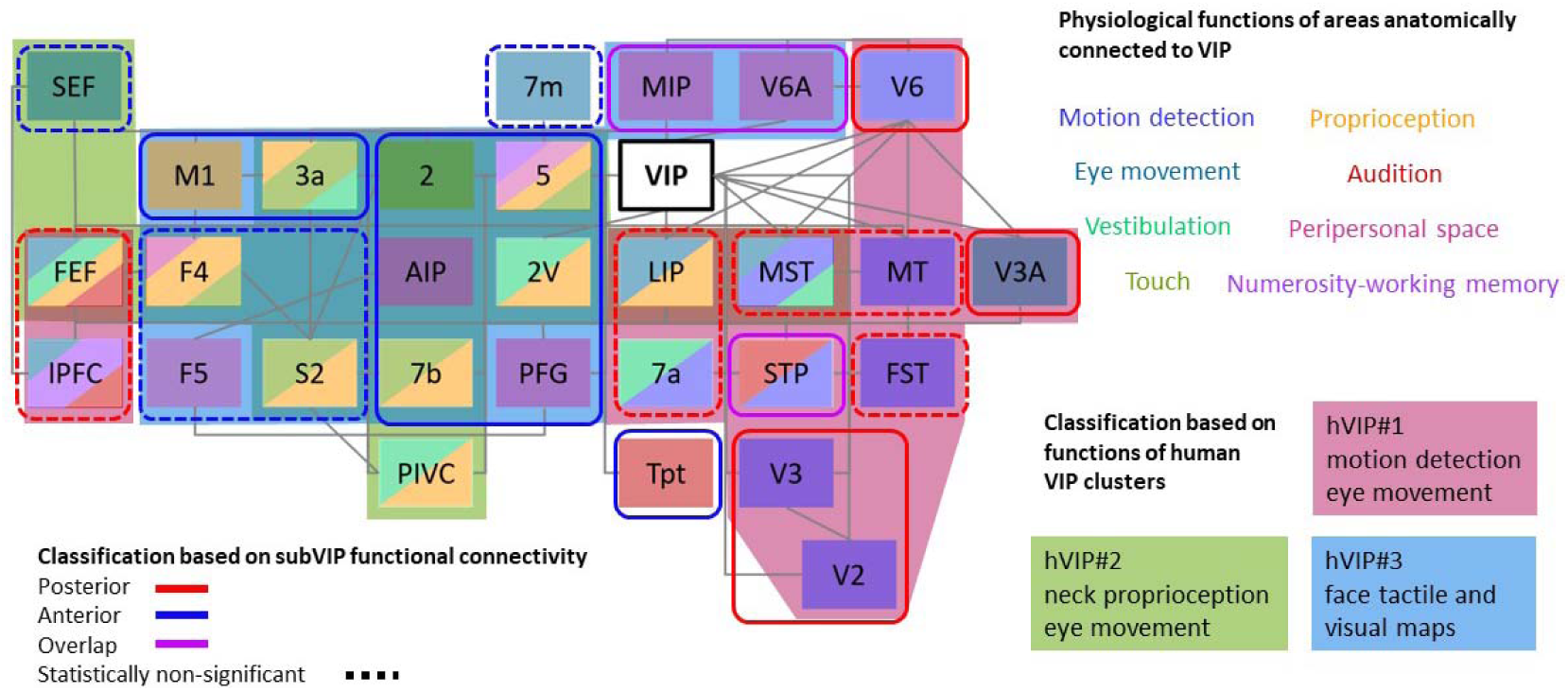
Macaque anterior and posterior ventral intraparietal area (VIP) functional connectivity match the functional specialization of human VIP clusters. Selected areas anatomically linked to VIP are arranged by relative anatomical locations, and text boxes are colored according to eight functional categories: visual motion, eye movement, vestibulation, touch, proprioception, audition, peripersonal space and numerosity/working memory. Striped text box colors show the overlap of multiple functional categories in the given area. Selected areas are classified based on matched functions relative to the three human VIP clusters (Foster et al., 2022). Background areas outside of the test boxes are colored according to hVIP#1 in pink, hVIP#2 in green and hVIP#3 in cyan. These colored areas are transparent to show overlap, for example, large overlap between hVIP#2 and hVIP#3 (green and blue). Selected areas are also classified based on their functional connectivity with either the anterior or posterior VIP ROIs as derived from the present study. Areas correlated to the posterior and the anterior VIP are circled in red and in blue solid lines, respectively, while areas correlated strongly to both VIPs are in purple. When the difference between the ratios of surface voxels correlated to the three VIP ROIs is not significant, areas are circled in dotted lines. If an area is statistically significant in only one hemisphere, we judge the significance based on the average z-score (supplementary Fig. 2). PFG is the rostral part of 7a. 7m: medial part of area 7, AIP: anterior intraparietal area, FEF: frontal eye field, FST: fundus of the superior temporal sulcus area, LIP: lateral intraparietal area, lPFC: lateral prefrontal cortex, M1: primary motor cortex, MIP: medial intraparietal area, MST: medial superior temporal area, MT: middle temporal area, PIVC: parieto-insular vestibular cortex, S2: secondary somatosensory area, SEF: supplementary eye field (F7), STP: superior temporal polysensory area, Tpt: temporoparietal area, V2: secondary visual area, V3: third visual area, VIP: ventral intraparietal area.

Macaque aVIP is functionally connected to the supplementary eye field (SEF), primary motor cortex (M1), F4, F5, 3a, 2, 2V, 5, 7m, AIP, secondary somatosensory cortex (S2), 7b, PFG and temporoparietal area (Tpt) (Fig. 6, circled in blue). This network is specialized in the processing of eye movements, vestibulation, touch, proprioception, audition and peripersonal space. It exhibits strong overlap with the functional specialization of human hVIP#2 and hVIP#3 for eye movement, tactile, proprioception and peripersonal space. However, because tactile and proprioceptive information are often processed in the same areas, it is difficult to associate anterior macaque VIP selectively with either hVIP#2 or hVIP#3. In contrast, areas that contribute to the processing of peripersonal space functionally segregate hVIP#2 from hVIP#3, as only hVIP#3 is associated with this function. These areas include F5, AIP and PFG that belong to the anterior VIP network. These areas also include V6A and the middle intraparietal area (MIP), both of which are strongly connected to both anterior and posterior VIP ROIs, possibly because they are very near to the seeds. Taken together, this points towards a functional homology between macaque anterior VIP and hVIP#3, and to a lesser extent hVIP#2.

Macaque pVIP is functionally connected to V2, V3, V3A, V6, medial temporal area (MT), medial superior temporal area (MST), fundus of the superior temporal sulcus area (FST), LIP, 7a, frontal eye field (FEF) and lateral prefrontal cortex (lPFC) (Fig. 6, circled in red). Areas in this network contribute mainly to motion detection and eye movements, which match with the functional specialization of human hVIP#1.

In sum, the functional specialization of the macaque anterior VIP functional connectivity network matches that of the lateral and the anterior human homologue clusters, while the posterior VIP functional connectivity network matches the posterior human homologue cluster. These results are in line with our hypothesis that functional specializations also exist within the macaque VIP along the anterior-posterior axis, and that this specialization within area macaque VIP can be viewed as a precursor of the more obvious separation into three distinct areas in the human brain. Tactile and proprioceptive functions are encoded in macaque aVIP, whereas motion detection and eye movement related functions are located in macaque pVIP. These connectivity-based results confirm prior observations about functional specialization within macaque VIP that were based on functional localizers in awake macaques (Guipponi et al., 2013, 2015).

In the present study, anterior, middle and posterior ROIs were composed of equivalent numbers of voxels. pVIP appears to uniquely match human hVIP#1, whereas anterior macaque VIP matches both human hVIP#2 and hVIP#3, while macaque mVIP may not have distinctive properties relative to macaque aVIP and pVIP. This might suggest a stronger functional specialization and expansion of middle and anterior relative to posterior human VIP, in agreement with the fact that cortical expansion in humans relative to macaques is more marked around the temporo-parietal junction (TPJ) and the inferior PPC than in medial PPC, that is, the areas in which these human VIP homologues are located (Van Essen and Dierker, 2007; Hill et al., 2010; Chaplin et al., 2013; Mantini et al., 2013). This match of cortical expansion with the emergence of hVIP#2 and hVIP#3 may also be why there is predominant colocalization of higher order cognitive functions in hVIP#2 and hVIP#3 relative to hVIP#1, for instance for peripersonal space representation and embodiment (Foster et al., 2022).

### The broader VIP functional connectivity networks serve social and cognitive functions in macaque

VIP functional connectivity networks include both areas that have been described to be anatomically connected to area VIP, and areas that are activated through indirect functional connectivity. We refer to these latter areas as the broader VIP functional connectivity networks. The macaque peripersonal space network includes four sub-networks (Foster et al., 2022; Cléry et al., 2015): MIP/V6A for reaching, 7b/F5/AIP for grasping, F4/VIP for self-defense and FEF/LIP for oculomotor control (Fig. 6). In addition, recent research demonstrated a strong social component in the functionality of peripersonal space (Cléry et al., 2015, 2017; Cléry and Ben Hamed, 2018, 2020; Serino, 2019), suggesting that the neural mechanisms underlying body ownership and peripersonal space are linked (Makin et al., 2008; Blanke, 2012; Blanke et al., 2015), and potentially also contribute to social processing (Orban et al., 2021). Although mirror self-recognition is generally not considered to be an intrinsic cognitive ability in monkeys (Gallup, 1982), it has recently been shown that macaques are capable of learning precise visual-proprioceptive associations of mirror images (Chang et al., 2017). Accumulating evidence in secondary somatosensory area S2 (Bretas et al., 2021) and in the insula (Schneider et al., 1993; Evrard et al., 2012) address the neural basis for body ownership in macaques. Interestingly, while S2 is anatomically connected to VIP, the insula is only functionally connected to VIP (suggesting an indirect anatomical connection). VIP could thus play an important role in body ownership of macaques, similarly to what has been described in the human hVIP#3.

In addition to the insula, there are other areas in the limbic, cingulate, orbitofrontal and prefrontal regions that are functionally, but not anatomically, connected to VIP. In the limbic regions, gustatory cortex (Kaskan et al., 2019) adds one more sensory modality to the multisensory properties of VIP, that has not been investigated to date. The temporal pole was found to contribute to the processing of familiar faces (Landi et al., 2021), while orbitofrontal area 12 processes information related to faces and emotion (Murphy and Bachevalier, 2020), as well as reversal learning (Rudebeck and Murray, 2011; Sallet et al., 2020). Related to this function, anterior cingulate cortex (Vogt et al., 1987) in the cingulate regions codes values and uncertainty needed for action selection (Monosov et al., 2020) via a network including orbitofrontal areas 11, 13 (Burke et al., 2014; Murray et al., 2015) and prefrontal areas 9, 46, 9/46 (Oemisch et al., 2015). More specifically, only area 24c (anterior cingulate sulcus) within area 24 is part of the broader VIP functional connectivity network, which signals reward allocation including during cooperative behavior (Chang et al., 2013). Partially overlapping with the value network, prefrontal area 44 controls vocalization (Petrides et al., 2005; Korponay et al., 2020) together with anterior/mid cingulate cortices and pre-supplementary/supplementary motor areas (Loh et al., 2017). Together, these considerations point towards a putative role of area VIP in a diverse set of higher order social and cognitive functions that is yet to be explored.

### Value of the study and implication for future experiments

The strength of the dataset on which we based our study is the high number of animals (n=10) and the fact that the investigated animals were awake during recording. The anesthetized state mostly reduces functional connectivity (Vincent et al., 2007; Grandjean et al., 2014; Hutchison et al., 2014; Thomas et al., 2021), while at the same time, some specific networks like visual and default networks actually show anesthesia-related increases in connectivity (Xu et al., 2018). Thus, awake state measurements provide a better understanding of the physiological functional networks.

The high number of individuals involved in this study enables group-level statistics for subareal ROI to whole-brain single-voxel connectivity analysis. The high spatial resolution functional connectivity reveals details within regions, for example, VIP-M1 connections concentrates laterally in M1 (Fig. 4) which corresponds to VIP’s specific role in face/head processing. Furthermore, detailed connectivity patterns also show correspondence across areas, like the medial and lateral branches of the pVIP to pVIP-mVIP gradient in the occipital regions. These functional connection details could guide future experiments that aim for parallel recordings across VIP and other areas to probe their inter-areal communications (Bastos et al., 2015; Dann et al., 2016; Semedo et al., 2019; Schneider et al., 2021). On the other hand, the newly identified broader VIP functional connectivity network sheds light on higher cognitive and social networks (Fujii et al., 2008; Jezzini et al., 2015; Serino, 2019; Villa et al., 2022), in which the role of VIP is yet to be explored.

Finally, the resting-state functional connectivity serves as a common research method for cross-species comparison between human and macaques (Gorges et al., 2017; Barron et al., 2020; Xu et al., 2020; Garin et al., 2022), as well as other non-human primates like marmosets (Hori et al., 2021; Garin et al., 2022). The abundant cytoarchitecture (Lewis and Van Essen, 2000a) and structural connectivity (Lewis and Van Essen, 2000b) data in macaque PPC compensates the incomplete data in human (Amunts et al., 2020; Niu et al., 2020) by assuming homology between the two species. Conversely, the rich fMRI studies in humans and the possibility to perform complex cognitive tasks also impact the understanding of brain functions and design of future experiments in non-human primates. Specific to this study, the three homologue human VIP clusters based on fMRI findings motivated the search of macaque VIP functional specialization based on intra-areal differential functional connectivity. Future research in humans can also probe the functional connectivity between the three hVIP clusters and the rest of the cortex for more detailed cross-species comparison, while in macaque, electrophysiology recordings along the anterior-posterior axis within IPS can help delineate the functional specializations at the neuronal level.

### Limitations of the dataset

Our finding that the functions of macaque VIP subdivision functional connectivity networks match the functional specializations of human homologue VIP clusters must be viewed in the light of some technical limitations. This study is uniquely based on the analysis of awake monkey resting state data in combination with an atlas-based approach by selecting voxels along the IPS fundus that covered VIP from the Markov map (Fig. 1). Confronting the observed functional connectivity between VIP and the rest of the brain with standardized and validated functional localizers would be very valuable to confirm the proposed functional gradient. Indeed, such localizers are expected to capture functional properties more consistently and precisely than atlas-based anatomical localization (Russ et al., 2021), especially when individual variation in VIP functional activations have been reported (Guipponi et al., 2013), as it has been shown that subject-specific ROIs are important for an enhanced functional resolution (Nieto-Castañón and Fedorenko, 2012). However, designing valid localizers for areas involved in multisensory and higher cognitive functions could be more challenging than for areas with low-level functional properties, such as primary somatosensory area S1 (Thomas et al., 2021). In addition, while the present study includes ten monkeys and, thus, provides evidence of cross-subject reproducibility of the observations, running functional localizers on such a large dataset is challenging in any given lab and will require concerted effort from multiple labs worldwide for several years (Milham et al., 2018, 2020, 2022).

## Data and code availability

Data and code are available upon request.

## Author contribution

Conceptualization: W-A.S. and S.B.H.; Data Curation: W.V., W-A.S., M.F., Formal Analysis: W-A.S. and S.B.H.; Funding Acquisition: S.B.H.; Investigation: W-A.S., and S.B.H.; Methodology: W-A.S., S.C., M.F. and S.B.H.; Resources: W.V.; Supervision: S.B.H.; Validation: S.B.H.; Visualisation: W-A.S., C.S.; Writing-Original draft: W-A.S. and S.B.H.; Writing -review & editing: W-A.S., S.C., M.F., W.V., T.H. and S.B.H.

## Ethical statement

Animal care and experimental procedures were performed in accordance with the National Institute of Health’s Guide for the Care and Use of Laboratory Animal, the European legislation (Directive 2010/63/EU) and were approved by the Animal Ethics Committee of the KU Leuven. Weatherall reports were used as reference for animal housing and handling. All animals were group-housed in cages sized 16-32 m^3^, which encourages social interactions and locomotor behavior. The environment was enriched by foraging devices and toys. The animals were fed daily with standard primate chow supplemented with fruits, vegetables, bread, peanuts, cashew nuts, raisins, and dry apricots. The animals were exposed to natural light and additional artificial light for 12 h every day. On training and experimental days, the animals were allowed unlimited access to fluid through their performance during the experiments. Using operant conditioning techniques with positive reinforcers, the animals received fluid rewards for every correctly performed trial. During non-working days, they received water in their living quarters. Throughout the study, the animals’ psychological and veterinary welfare was monitored daily by the veterinarians, the animal facility staff, and the lab’s scientists, all specialized in working with non-human primates. The animals were healthy at the conclusion of our study. M1 and M2 are currently still employed in other studies.

## Acknowledgments

We thank also thank Thomas Perret, Johan Pacquit and Marco Bimbi for help with the hardware computational resources.

## Supplemental data

**Supplementary Fig. S1.**
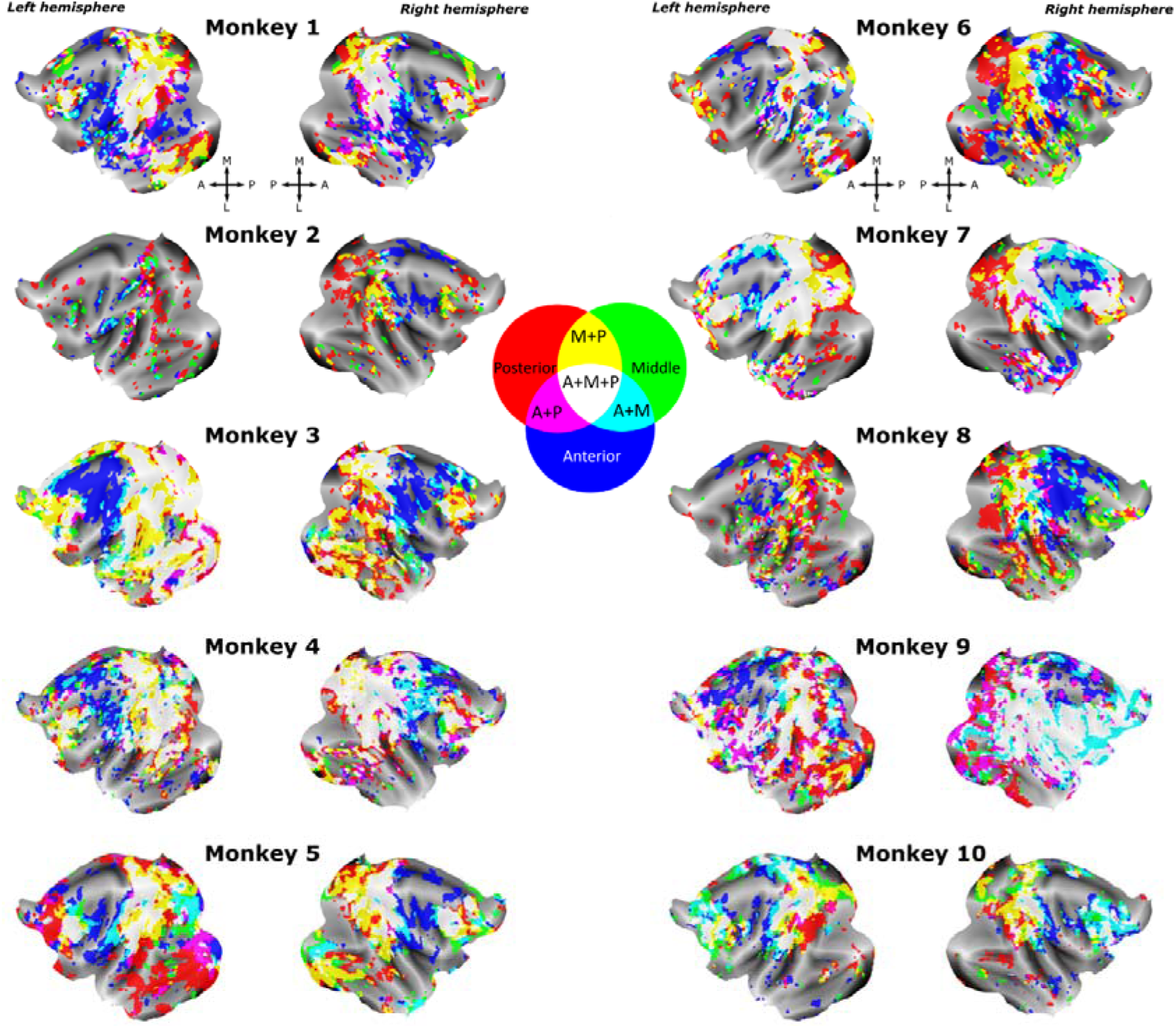
Topographical organization of the cortical regions that are uniquely functionally connected to one of the aVIP, mVIP and pVIP ROIs, or to two or more of these three ROIs in each of the ten individual monkeys. Flatmaps of both hemispheres show unique and overlapping regions for the three VIP connectivity maps. Surface voxels with z-scores above 0.05 are colored. Unique, non-overlapping voxels functionally connected to the anterior, middle and posterior VIPs are colored in blue, green and red, respectively. The anterior-middle, middle-posterior and anterior-posterior overlapping voxels are coded in cyan, yellow and magenta. Regions functionally connected to all three seeds are shown in white. AS, arcuate sulcus; CiS, cingulate sulcus; CS, central sulcus; IOS, inferior occipital sulcus; IPS, intraparietal sulcus; LS, lateral sulcus; LuS, lunate sulcus; OTS, occipitotemporal sulcus; PS, precentral sulcus; STS, superior temporal sulcus.

**Supplementary Fig. S2.**
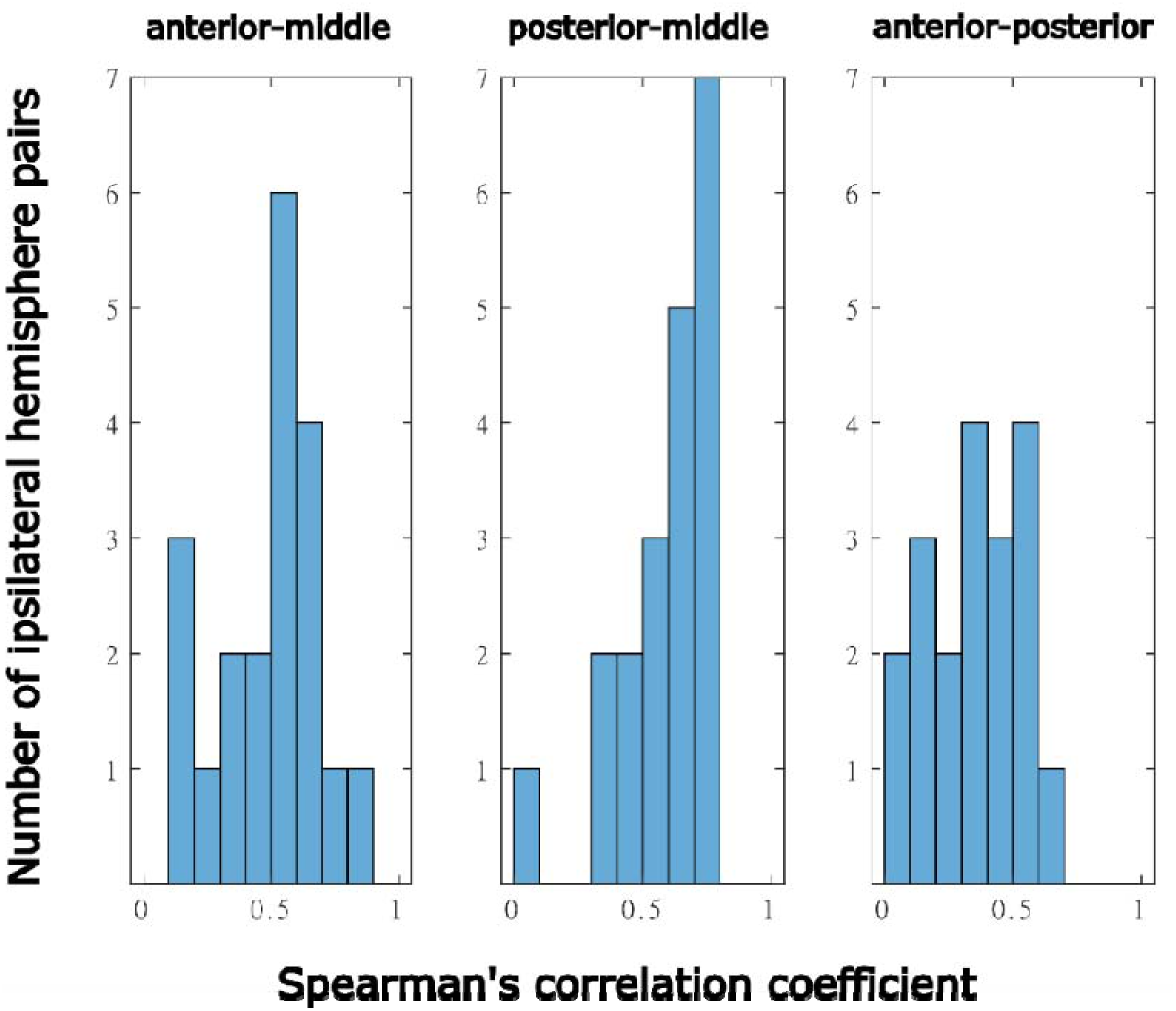
Distributions of spearman’s correlation coefficient between the ipsilateral aVIP, mVIP and pVIP whole brain connectivity maps of the 10 individual monkeys. Spearman’s correlation coefficient calculated between ipsilateral pairs of anterior-middle (aVIP-mVIP), posterior-middle (pVIP-mVIP) and anterior-posterior (aVIP-pVIP) VIP connectivity maps of each monkey, two hemispheres plotted together. All correlation coefficients have p values <0.05.

**Supplementary Fig. S3.**
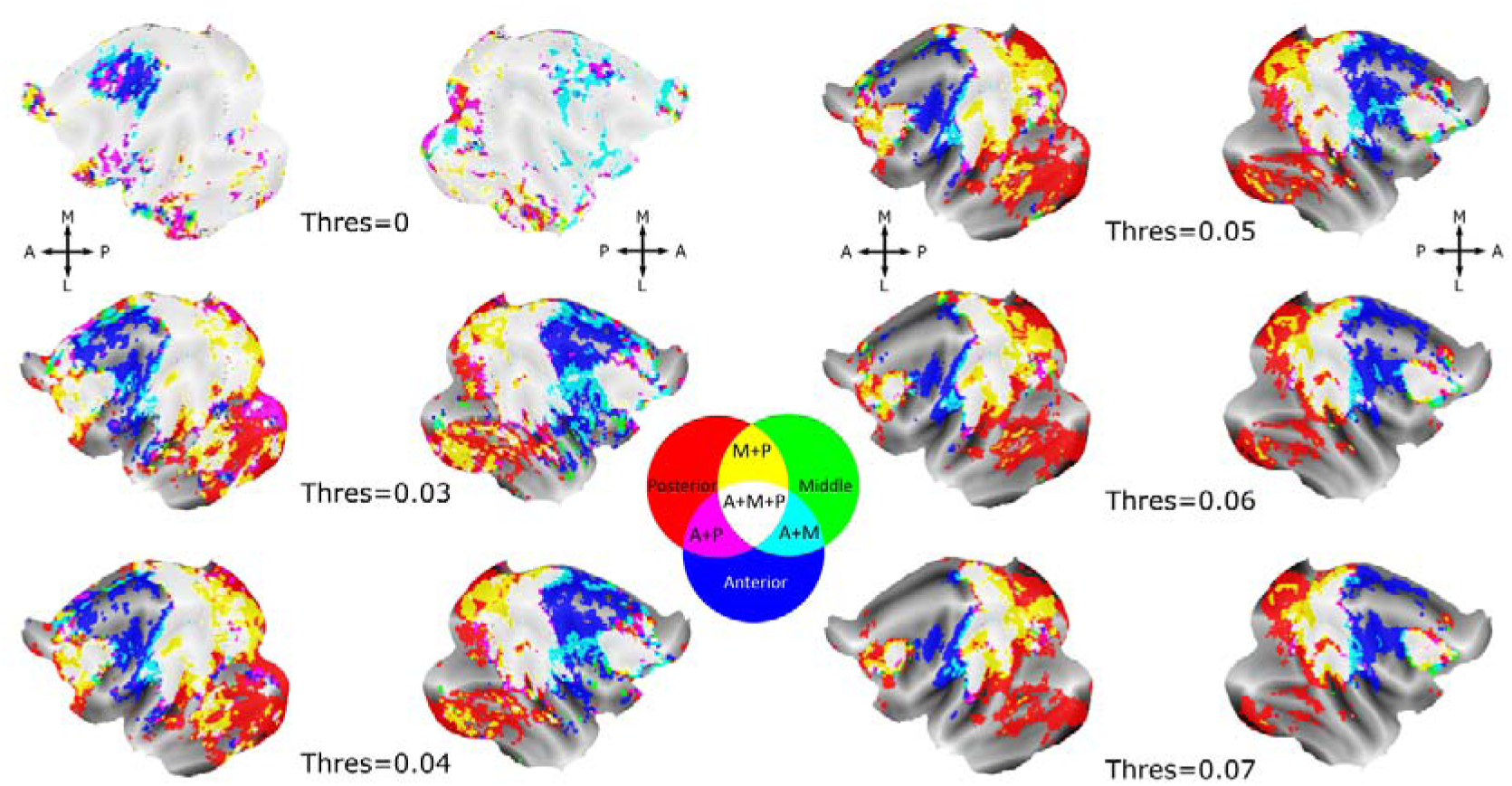
Different z-score thresholds to obtain topographical organization of the cortical regions that are uniquely functionally connected to one of aVIP, mVIP and pVIP ROIs, or to two or more of these three ROIs. Average flatmaps of both hemispheres show unique and overlapping regions for the three VIP connectivity map. Surface voxels with average z-scores above different thresholds (0 to 0.07) are colored. Unique, non-overlapping voxels functionally connected to the anterior, middle and posterior VIPs are colored in blue, green and red, respectively. The anterior-middle, middle-posterior and anterior-posterior overlapping voxels are in cyan, yellow and magenta. Regions functionally connected to all three seeds are in white. Pie charts show percentages of different categories of unique and overlapping surface voxels in the left and the right hemispheres with the same color code as in the flat maps. Black color in pie charts represents percentage of surface voxels without any signals.

**Supplementary Fig. S4.**
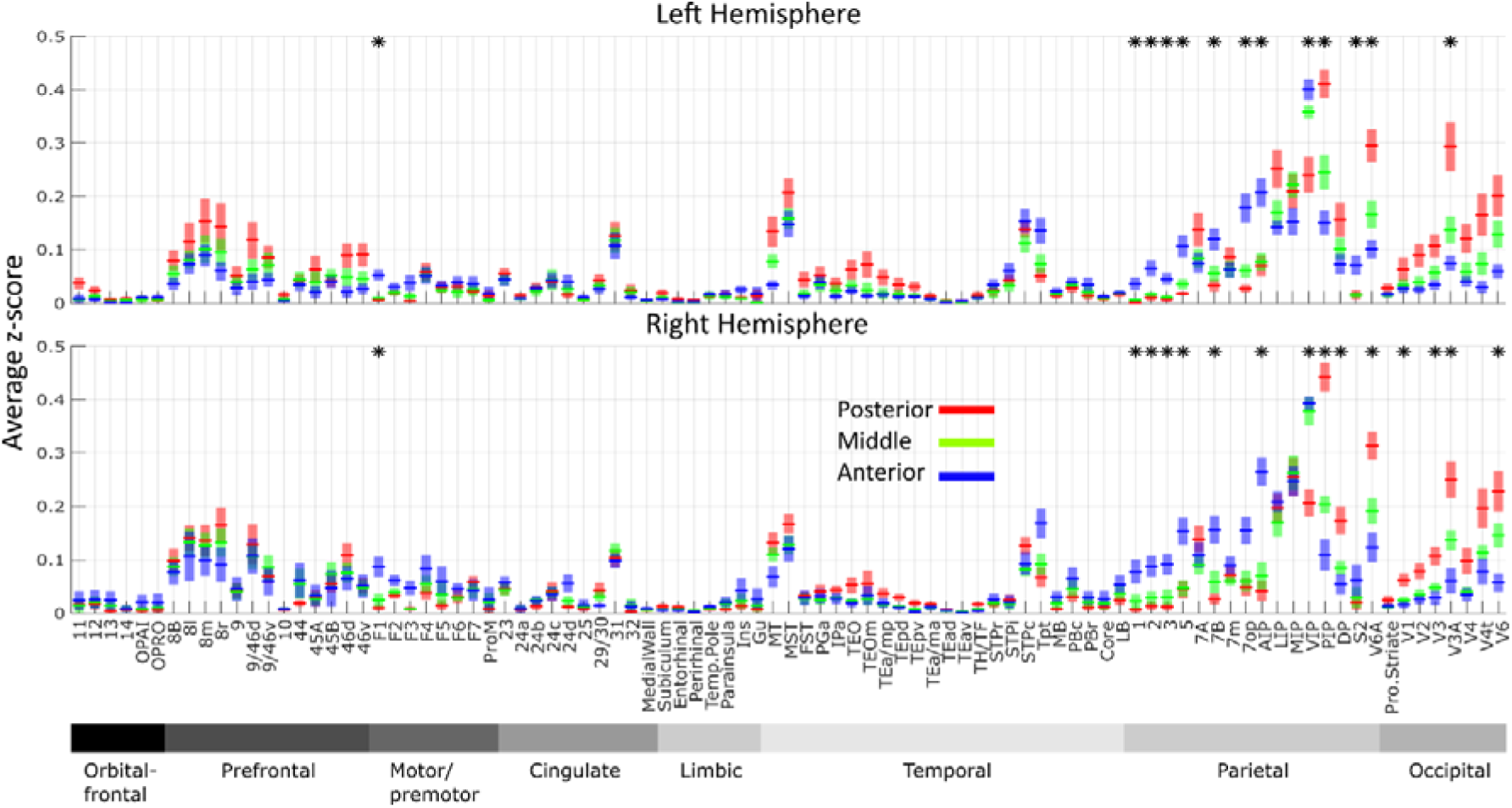
Average z-scores of functional correlation between aVIP, mVIP and pVIP in atlas-defined cortical functional regions and areas. For each area (Markov atlas, Markov et al., 2014), average z-scores of surface voxels correlated to aVIP (blue), mVIP (green) and pVIP (red) are calculated separately and plotted as line charts for the right and the left hemispheres separately. Shadows show standard error calculated across animals. Areas for which the average z-scores of surface voxels correlated to the three VIP seeds are significantly different (Kruskal-Wallis test with Benjamini-Hochberg procedure for multiple comparisons correction, p<0.05) are marked with a star on top. The areas are grouped by brain regions, in the following order: orbitofrontal, prefrontal, motor/premotor, cingulate, limbic, temporal, parietal and occipital. This arrangement allows most of the areas to be continuous in space with its previous and following areas in the figure.

